# The Brain Encodes Pure Logic Beyond Natural Language and At The Boundaries With Mathematics

**DOI:** 10.64898/2026.08.24.746607

**Authors:** Marie Amalric, Gabriele Pierguidi, Michela Terzani, Greta Gaimarri, Paula A. Maldonado Moscoso, Chiara Di Domenico, Giorgio Vallortigara, Simone Viganò, Manuela Piazza

## Abstract

You are not not reading this sentence. True, because you are. It’s the law of double negation, a logical rule that states that a statement that is not not true is itself true. The development of pure logic — the study of the properties of formal systems, irrespective of their content — represents one of the most abstract intellectual achievements of human culture. Yet its neural correlates remain vastly unexplored. Is pure logic represented in a linguistic format like natural language, in an abstract symbolic format like other formal systems such as mathematics, or as a fully distinct domain of knowledge? To address this question, we monitored the brain activity of professional logicians and matched controls while they evaluated the truth of spoken statements of pure logic, and compared these activations to those elicited by language and mental arithmetic. We show that pure logic recruits left-lateralized brain circuits that are completely distinct from those involved in language processing. Those circuits only partially overlap with those involved in mathematics. These findings highlight an independent neural pathway for abstract thought and indicate that the brain’s capacity to process structured rules of formal systems is implemented beyond language and at the boundaries with mathematics.

## Introduction

Humans are remarkable for their ability to think abstractly. Thanks to our capacity to use arbitrary symbols and formal systems to manipulate them, we can reason abstractly not only about objects, feelings, and events, but also about concepts, rules, and relations. Many domains of human knowledge rely on such symbolic systems: Arithmetic manipulates symbols denoting quantities by means of operators; algebra operates on variables and functional relations; linguistics formalizes the structure of natural language through grammars and syntactic rules; and propositional logic uses abstract operators such as conjunction, disjunction, and negation, together with rules of inference. Although these domains differ in content, they share a reliance on symbolic representations and structured transformations. Within this landscape, pure logic, a recent discipline that emerged thanks to the crucial contributions of philosophers/mathematicians such as Russell, Hilbert, and Gödel, occupies a very distinctive position^1^. Contrary to the other formal systems, pure logic studies the structure and properties of formal systems themselves, irrespective of their referents. It examines notions such as validity, consistency, completeness, definability, and computability, which are properties that can apply to any symbolic framework. For example, the notion of consistency (whereby a formal system is consistent if it cannot derive both a statement and its negation), applies equally to arithmetic and to propositional logic. Pure logic is therefore a “meta-language” that straddles natural language and mathematics, in that it borrows linguistic expressions to formulate propositions while borrowing mathematical formalisms for its rigorous analysis, raising the question of how such an ultra-abstract form of reasoning is implemented in the human brain.

A growing body of neuroimaging research has begun to characterize the regions of the brain that are involved in formal symbolic systems such as arithmetic, high-level mathematics, computer programming, and logical reasoning. Arithmetic, more advanced mathematics, and computer coding were all systematically shown to recruit a bilateral fronto-parietal network (including the IPS and MFG, and the pITG) that is almost completely dissociated from the language network^2–8^. Results on logical reasoning, primarily investigated with syllogistic reasoning tasks, are instead less consistent, sometimes linked to a left-lateralized fronto-parietal circuit dissociated from language, and sometimes with a left-lateralized circuit that overlaps with language^9–14^. To date, virtually nothing is known about how the brain represents the kind of objects and rules that are common across such different formal systems, namely pure logic.

Because most previous neuroscientific studies of logic have relied on tasks involving logical inferences (e.g., evaluating syllogisms or conditional inferences) and have been conducted in non-experts, our ability to capture the mental representation of pure logic as a domain of abstract knowledge remains extremely limited. Just as studying basic arithmetic cannot reveal how the brain represents algebra or topology, restricting the study of logic on syllogistic reasoning does not explain how the brain represents proof systems or model theory. To advance beyond this limitation, one must investigate experts who have acquired pure logic as a specific domain of knowledge.

In the present work, our main focus is on the neural representation of pure logic when processed in its natural linguistic form, that of verbally expressed theorems. Our main question is whether pure logic, a discipline that straddles natural language and mathematics, is represented in the brain through a linguistic code, through a mathematical code, or as fully separate from both language and mathematics. To answer this question, inspired by previous work on professional mathematicians^7^, we scanned professional logicians and matched controls while they evaluated the truth of spoken statements, matched in length and syntactic complexity, that either referred to advanced pure logic concepts or to general knowledge concepts (see SOM for the full list of statements presented, and Figure 1A). To further characterize the relation between logic, mathematics, language and visual processing, we also scanned subjects with additional fMRI runs evaluating their brain activity during sentence processing, mental arithmetic, and visual processing of faces, bodies, tools, houses, numbers, letters, and written logic expressions.

**Figure 1.**
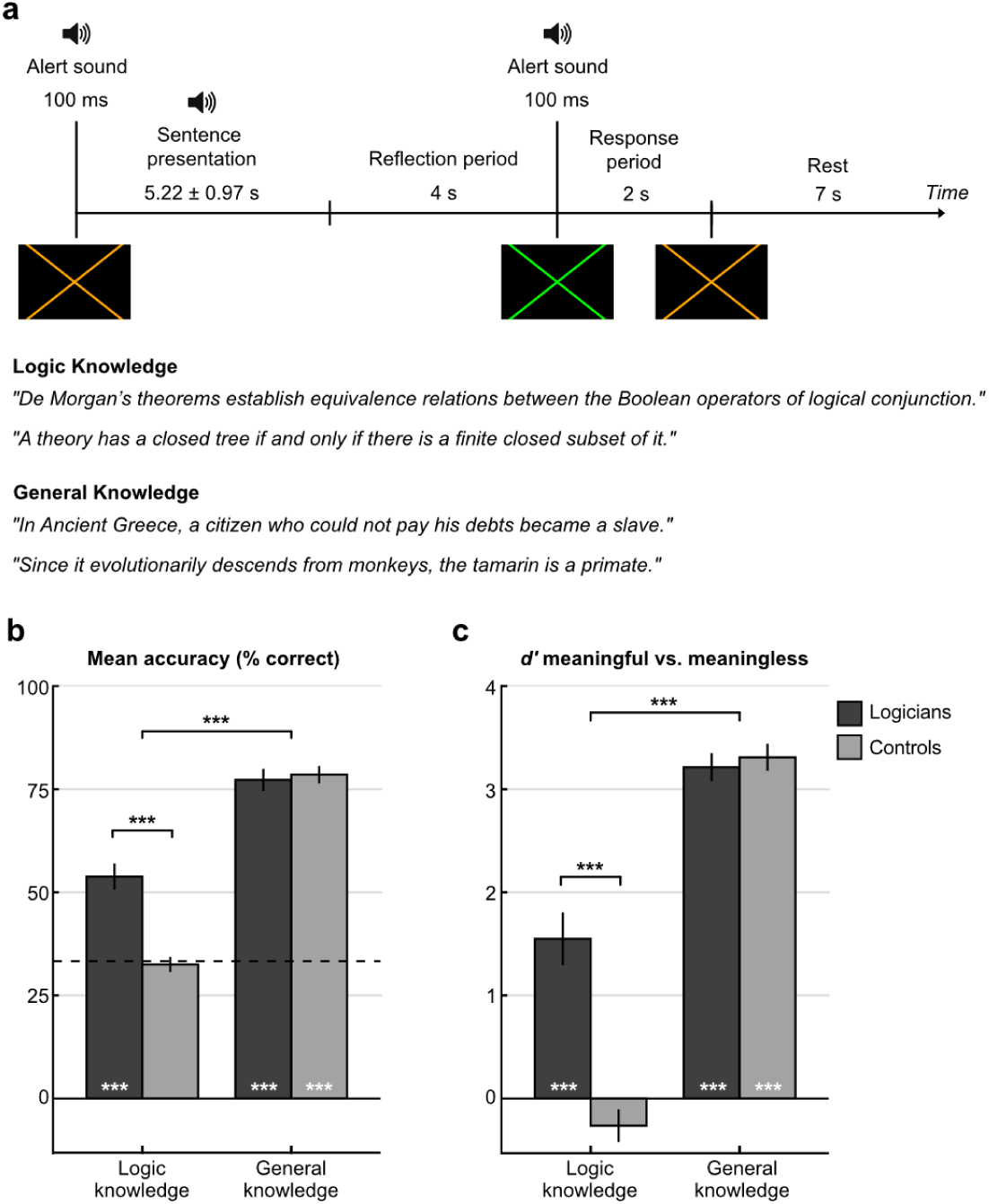
Main experimental paradigm and behavioral results. (a) Within each trial, participants listened to spoken statements and after 4 s they were asked to decide whether the statement was true, false, or meaningless via button press. (b) Task accuracy (% correct responses) for the two different statement categories and the two groups of participants. (c) Average d′ values reflecting discrimination between meaningful and meaningless statements. *** p < 0.001. Error bars represent the standard error of the mean (SEM).

## Results

### Behavioral results

#### Performance in the scanner: logicians and controls performed equally on general knowledge statements, but only logicians could evaluate the truth values of pure logic statements

Logicians performed above chance (33.3%) for both logic statements (54 ± 3.1%; t(11) = 6.5, p<0.001) and general knowledge statements (77 ± 2.7%; t(11) = 16.2, p < 0.001) (Figure 1B). In contrast, control participants performed above chance only for general knowledge statements (78 ± 2.1%; t(9)=21.46, p<0.001) but not for logic statements (32 ± 1.8%; t(9)=−0.43, p=0.67). A 2×2 mixed ANOVA with Condition as a within-subject factor and Group as a between-subject factor revealed significant main effects of Group (F(1,20) = 11.03, p < 0.005) and Condition (F(1,20) = 264.26, p < 0.001), as well as a significant Group × Condition interaction (F(1,20) = 27.97, p < 0.001). In particular, logicians and controls’ performance did not differ for general knowledge statements (t(20) = −0.36, p = 0.7) while they did so for logic statements (t(20) = 5.54, p < 0.001). To quantify participants’ discrimination between meaningful and meaningless statements, we then used signal-detection analysis and found converging results (Figure 1C). Logicians showed reliable discrimination between true and false statements for both logic (d′ = 1.55 ± 0.25; t(11) = 6.04, p < 0.001) and general knowledge statements (d′ = 3.2 ± 0.13; t(11) = 23.58, p < 0.001). In contrast, controls showed reliable discrimination only for general knowledge statements (d′ = 3.3 ± 0.13; t(9) = 25.35, p < 0.001), with no evidence of discrimination for logic statements (d′ = −0.26 ± 0.16; t(9) = −1.67, p = 0.13). A 2×2 mixed ANOVA confirmed significant main effects of Group (F(1,20) = 16.4, p < 0.005) and Condition (F(1,20) = 292.73, p < 0.001), as well as a significant Group × Condition interaction (F(1,20) = 38.84, p < 0.001). Between-group comparisons identified a significant difference for logic (t(20) = 5.72, p < 0.001) but not for general knowledge (t(20) = 0.50, p = 0.62). These results suggest that participants performed the task well above chance, and that only logicians reliably evaluated meaningful statements in both logic and general knowledge domains.

#### Post-MRI questionnaire: in logicians, general knowledge statements were found to be easier, better understood, and more confidently assessed compared to pure logic statements

We next analyzed the subjective variables rated in the post-MRI questionnaire (see Figure S1). Within the group of logicians, both logic and general knowledge statements were reported to be solved based on declarative knowledge retrieval more than on active computation (on a subjective scale converted to a 0-100 score; meaningful logic statements: 28.35 ± 4.72; meaningful general knowledge statements: 26.97 ± 3.84; t(11) = 0.21, p = 0.84). General knowledge statements, compared to meaningful logic statements were better understood (logic: 74.2 ± 3.87; general knowledge: 91.9 ± 1.75; t(11) = −4.13, p < 0.005), and rated less difficult (logic: 50.5 ± 4.24; general knowledge: 31.5 ± 4; t(11) = 5.21, p < 0.005). In assessing general knowledge statements, participants were also more confident (logic: 55.82 ± 3.77; general knowledge: 76.73 ± 2.39; t(11) = −3.8, p < 0.005), and reported higher mental imagery (logic: 9.86 ± 2.29; general knowledge: 17.67 ± 4.64; t(11) = −2.25, p = 0.046). Ratings for general knowledge statements did not differ between logicians and control participants for any of the subjective variables (all ps > 0.14). A binary logistic regression within the group of logicians revealed significant correlations between accuracy in the scanner and subsequent ratings of comprehension (r = 0.28, p < 0.001), confidence (r = 0.35, p < 0.001) and difficulty (r = −0.22, p < 0.001). Perhaps unsurprisingly, logicians’ ratings of comprehension were negatively correlated with difficulty ratings (r = −0.26, p < 0.001), indicating that higher levels of sentence comprehension were associated with lower perceived difficulty.

### fMRI results

#### Brain responses to pure logic statements: in logicians pure logic statements activate a highly left lateralized fronto-parieto-temporal network

To characterize the neural systems supporting the processing of pure logic content, we contrasted brain responses to meaningful statements involving logic concepts with those involving general knowledge in logicians. Logic statements elicited increased activity in a left-lateralized fronto-temporal-parietal network, including the left intraparietal sulcus (left IPS: [−50 −44 51], t = 7.88), the left middle frontal gyrus (left MFG: [−46 42 26], t = 5.59; all MNI_x,y,z_ coordinates), and bilateral posterior inferior temporal gyri (left pITG: [−53 −54 −16], t = 8.78; right pITG: [54 −47 −14], t = 5.84) (Figure 2A). Single-subject-based analyses of the lateralization indices (see Methods) confirmed that the logic-responsive network was highly left-lateralized (Lateralization Index (hereafter LI) = 0.48 ± 0.07; t(11) = 5.99, p < 0.001; Figure 2C). Plots of mean activation at the peaks in these regions revealed greater activity in logicians than controls for logic statements (Figure 2A). Importantly, this circuit was qualified by a significant interaction between Content (logic > general knowledge) and Group (logicians > controls), confirming that activation elicited by logic statements was higher in logicians than in controls, thus indicating that it was driven by the comprehension of the logic content, and not by their specific surface forms (Figure 2B).

**Figure. 2.**
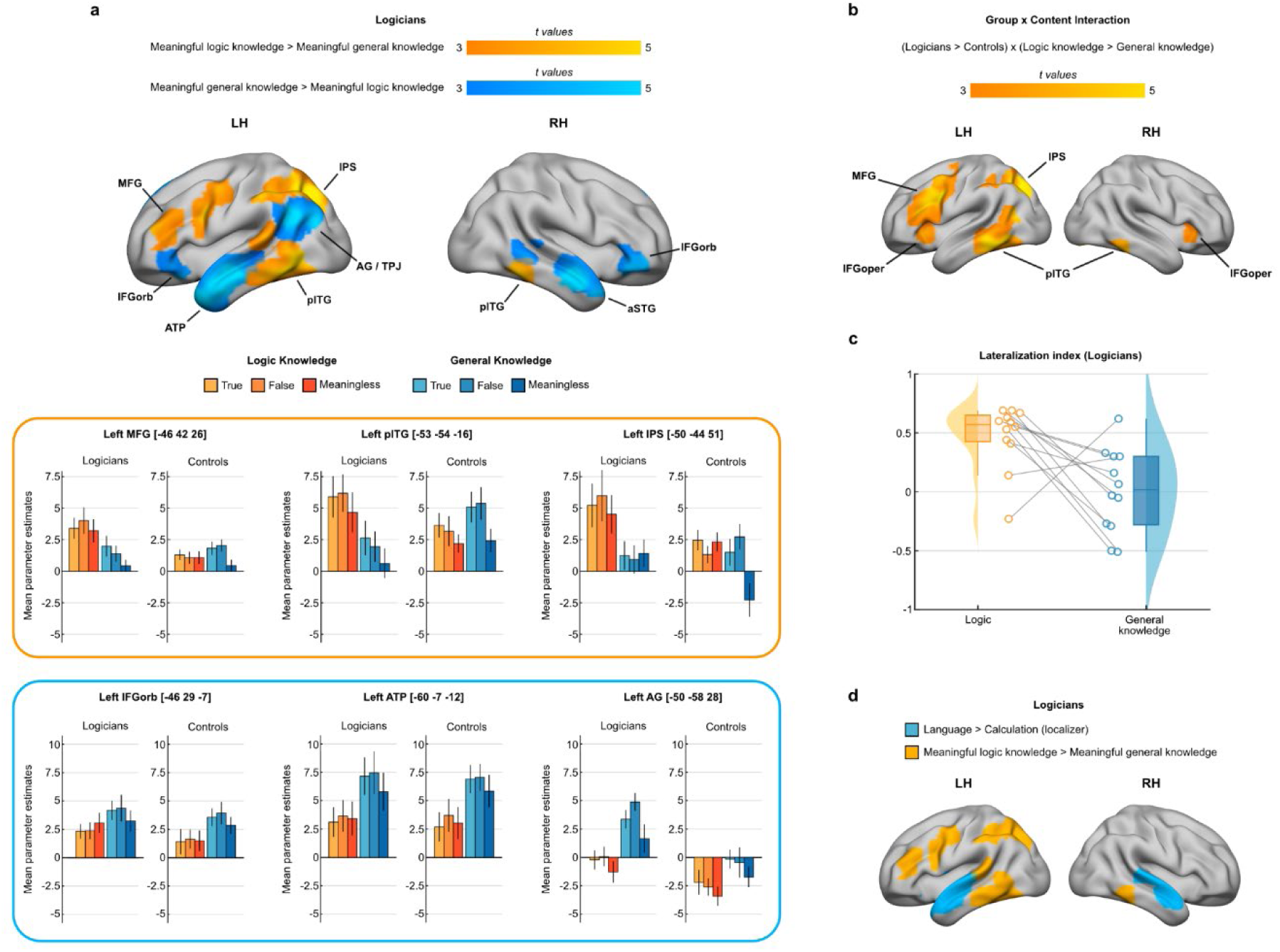
Distinct brain areas for pure logic expertise and for general knowledge in logicians. (a) Top: Whole-brain view of areas activated during processing of meaningful logic statements (yellow) versus meaningful general knowledge statements (blue). In this and all subsequent figures, brain maps are thresholded at voxel p < 0.001, cluster at pFDR < 0.05, corrected for multiple comparisons across the brain volume. Bottom: Mean activity estimates for each statement category extracted from the main activation peaks for logic (orange) and general knowledge (blue). In this and all subsequent figures, error bars represent standard error. (b) Group x content interaction map indicating a greater difference between meaningful logic and general knowledge statements in logicians than in controls. (c) Lateralization index of the circuits involved in logic and general knowledge. (d) The logic expertise network (yellow) and the language areas (blue) derived from the language localizer (contrast sentences > mental calculation).

#### Segregation between pure logic and language: general knowledge statements activate the classical language areas: bilateral superior temporal gyrus, angular gyrus, and inferior frontal gyrus

In contrast, general knowledge statements recruited regions in the temporal poles and anterior superior temporal gyri (left TP/aSTG: [−60 −7 −12], t = 11.12; right TP/aSTG: [−64 −4 −16], t = 11.25), angular gyri at the temporoparietal junction (left AG/TPJ: [−50 −58 28], t = 9.08), and the orbital portion of the inferior frontal gyrus of both hemispheres (left IFGorb: [−46 28 −7], t = 5.43; right IFGorb: [43 39 −9], t = 6.54; Figure 2A). In these regions, plots of mean activation showed the opposite pattern to that observed in the logic-responsive network: greater activity for general knowledge than for logic statements, minimal or no response to logic statements, and no significant group difference for meaningful general knowledge statements (except in AG). Single-subject-based analysis of lateralization index confirmed a bilateral engagement for general knowledge processing (LI = 0.01 ± 0.1, t(11) = 0.1, p = 0.92, Figure 2C), which was significantly more bilateral compared to that of the logic-responsive network (t(11) = −2.96, p = 0.013, Figure 2C).

#### The segregation between pure logic and language is confirmed with an independent language localizer

In order to further explore the dissociation between the logic-responsive network and the language network in logicians we used an independent language comprehension localizer, adapted from Pinel and collaborators^15^, in which participants processed simple spoken or written sentences and arithmetic calculation problems. Within this localizer, the contrast *sentences > calculation* identified the bilateral superior temporal sulcus/gyrus and anterior temporal cortex. While group-level maps revealed substantial spatial overlap between the language localizer and general knowledge-related activations in the main task, the language regions identified by the localizer showed no overlap with those activated by logic statements (Figure 2D). This segregation was confirmed by a sensitive single-subject quantification of whole-brain overlap between logic-responsive voxels and the localizer language network (Dice coefficient, a value that indicates the proportion of shared area across the single participants’ activity maps, see Methods). This overlap was minimal (Dice = 0.009 ± 0.006; no participant with significant overlap in hypergeometric tests; Z = −14.88; p = 0.99) and significantly lower than the overlap between general knowledge (main experiment) and language (localizer) regions (Dice = 0.084 ± 0.023; t(10) = −2.98, p = 0.014; 10 participants with p < 0.01 in hypergeometric tests; Z = 22.34; p < 0.001). These findings further confirm that despite being expressed in matched linguistic formats, logic and general knowledge rely on dissociated brain circuits.

#### The dissociation between pure logic and language is not determined by differences in task difficulty

To verify that the different networks identified for pure logic and for general knowledge in logicians were not the result of a different level of difficulty associated with the two conditions (see above, “Performance in the scanner”, and “Post-MRI questionnaire” paragraphs), we ran several control analyses. First, we repeated the whole-brain analyses while including participants’ mean accuracy as a covariate of objective difficulty at the second level. Average accuracy in the scanner was used instead of subjective ratings from the post-MRI questionnaire to avoid uncontrolled inter-individual differences in the use of the scale in the questionnaire.

This analysis revealed that logic content continued to engage the same left-lateralized fronto-parietal-temporal network, including the IPS, MFG, and pITG, as observed in the main analysis (Figure 3A). Second, we performed a within-subject analysis based on subjective difficulty ratings in the post-MRI questionnaire, contrasting logic statements rated as easy with general knowledge statements rated as difficult. Easy vs. difficult trials were determined on the basis of a subject-specific mean-split over their difficulty ratings, a method that is not impacted by inter-individual differences in the use of the scale. As a result, easy logic statements were evaluated as being easier compared to difficult general knowledge statements (easy logic statements = 29.41 ± 4.78; difficult general knowledge statements = 49.77 ± 4.52; difference t(11) = −3.87, p < 0.005, Figure 3B). Despite the reversal in subjective difficulty, comparing the brain activation maps for easy logic vs. difficult general knowledge statements revealed the same map found initially, when we analyzed all statements (easy and difficult) together: greater activation in the Left IPS, MFG, and pITG, (Figure 3C and 3D). These two control analyses prove convergently that the logic-related activations identified in the main contrasts do not depend on differences in task difficulty or cognitive effort but instead reflect the processing of pure logic content itself.

**Figure 3.**
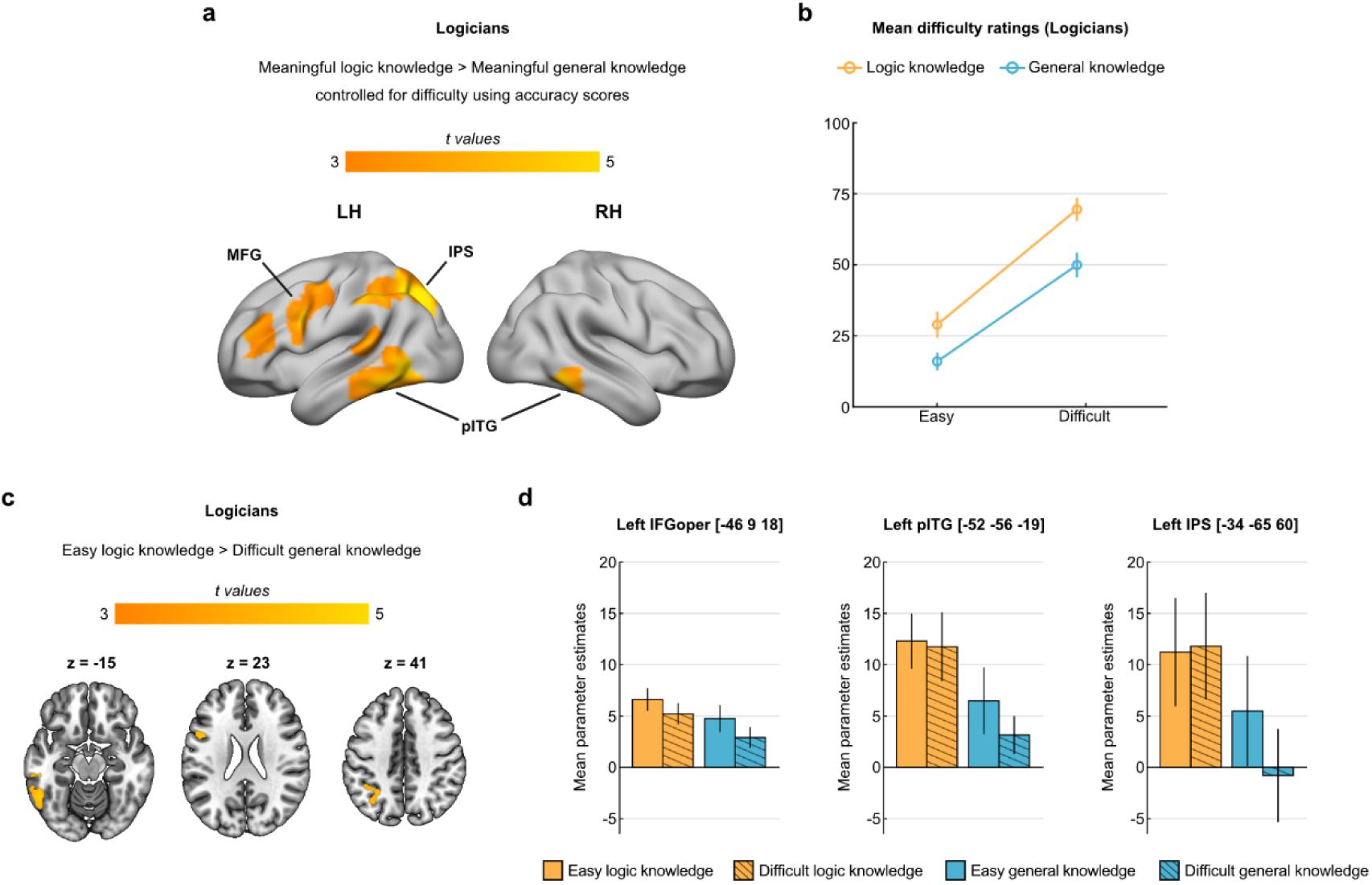
Controls for task difficulty (a) Results of the contrast logic > general knowledge from a GLM model where accuracy was entered as a covariate regressor. (b) Mean difficulty ratings from the post-MRI questionnaire for easy and difficult meaningful logic and general knowledge statements. (c) Axial slices showing the main regions activated in logicians by the contrast [easy logic > difficult non-logic] across all meaningful statements. (d) Mean activity estimates for easy and difficult meaningful logic and general knowledge statements extracted from the main activation peaks in (c).

#### The pure logic network partially overlaps with the calculation network

A qualitative inspection of the brain regions activated by logic statements in logicians revealed a marked similarity with those previously reported for mathematical statements in mathematicians ^7,8^ (Figure 4A), with the crucial exception of right hemispheric activations, that were almost entirely absent for pure logic. Although in the current experiment we did not include mathematical statements, thanks to our independent localizer, we did have the opportunity to isolate, on an individual subject basis, the brain networks involved in arithmetic calculation, which was previously shown to substantially overlap with that of advanced mathematics^7^. Thus, to statistically assess the correspondence between logic-related and math-related activations, we evaluated the spatial overlap of pure logic-related activations with brain responses to calculation, identified using the contrast *calculation > sentences* of our independent localizer (Figure 4B). Single-subject Dice analysis demonstrated a substantial overlap between the two networks, which was statistically significant in both hemispheres (Left hemisphere Dice = 0.16 ± 0.03; 10 participants with p < 0.01 in hypergeometric tests; Z = 22.05; p < 0.001; Right hemisphere Dice = 0.06 ± 0.019; 9 participants with p < 0.01 in hypergeometric tests; Z = 17.95, p < 0.001), but much more predominant in the left hemisphere (Left vs. Right Logit-transformed Dice paired t-test: t(10) = 2.67, p = 0.025). The calculation network itself showed a more bilateral profile of activation (LI = 0.18 ± 0.1, t(10) = 1.82, p = 0.098; Figure 4C). In sum, in the left hemisphere logic largely overlaps with the calculation network (left pITG, left parietal cortex, left middle frontal gyrus and left dorsolateral prefrontal cortex) and in the right hemisphere only with the pITG component.

**Figure 4.**
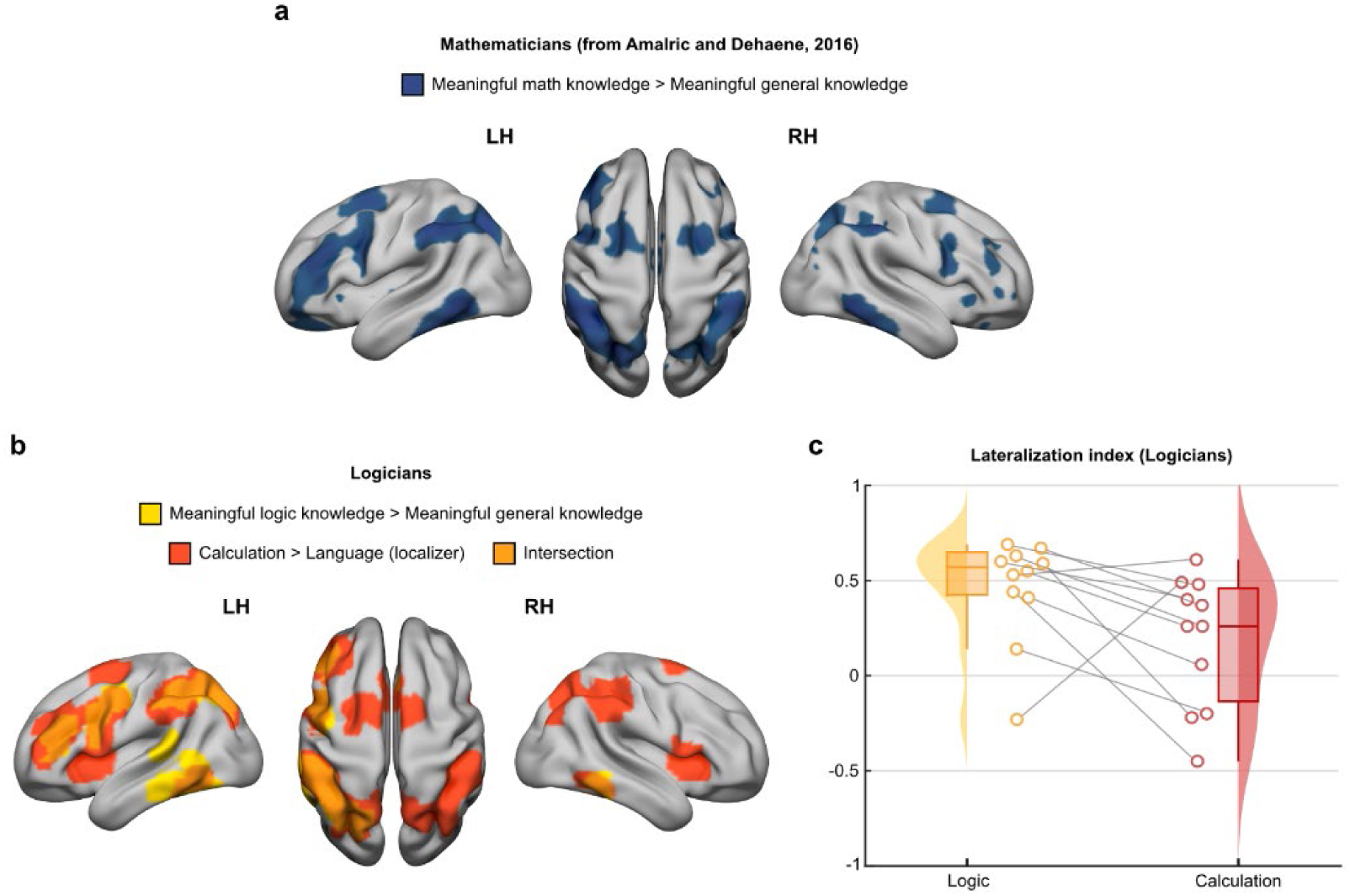
(a) The math-related network in mathematicians, from Amalric and Dehaene, 2016. (b) The logic-related network (yellow) and the mental arithmetic network (red) derived from the localizer (contrast mental calculation > sentences) and their intersection (orange), in logicians. (c) Lateralization index of the circuits involved in logic and calculation.

#### The representations of written logic formulae in the brain

Prior work has shown that, in adult skilled readers, written symbols (words and numbers) recruit category selective territories of the ventral visual stream, which are adjacent to, but separated from, those responding to other visual categories^16–19^, and that expertise often generates a reorganization of the cortical territories dedicated to visual processing^7,20,21^. Here we examined whether written logic formulae elicit ventral stream response distinct from or overlapping with those evoked by other visual stimuli, in particular other visual symbols such as words and numbers, and whether expertise changes the ventral stream response to other categories of visual stimuli. To answer these questions, at the end of the fMRI session, we presented, in a separate run, written logic formulae (strings composed of standard logical operators and symbols arranged as formal expressions) along with images pertaining to six other categories: faces, houses, tools, numbers, words, and checkerboards (see Figure 5a for some examples of stimuli). Results indicate that the typical mosaic of ventral occipito-temporal preferences for the different categories (obtained by contrasting the activity for each of the categories vs. all the others, excluding formulae), did not differ in logicians and controls (no significant voxel resulted from the interaction between visual category and group). Moreover, they revealed that in both logicians and controls formulae activated a set of voxels in the bilateral pITG and IPS more than all other visual stimuli. The intensity of this formula-selective activity did not differ across groups, as evident from whole-brain group comparisons (interaction group x category, yielding no significant voxel) as well as peak and ROI analyses (interaction group x category; both ps > 0.5). However, its extent tended to differ across groups, particularly in the right pITG compared to the left pITG, where written formulae evoked a broader patch in logicians compared to controls (interaction Group x Hemisphere: F(1,20) = 4.28, p = 0.052, t-test in the right hemisphere: t(20) = 1.82, p = 0.084, Cohen’s d = 0.81).

**Figure 5.**
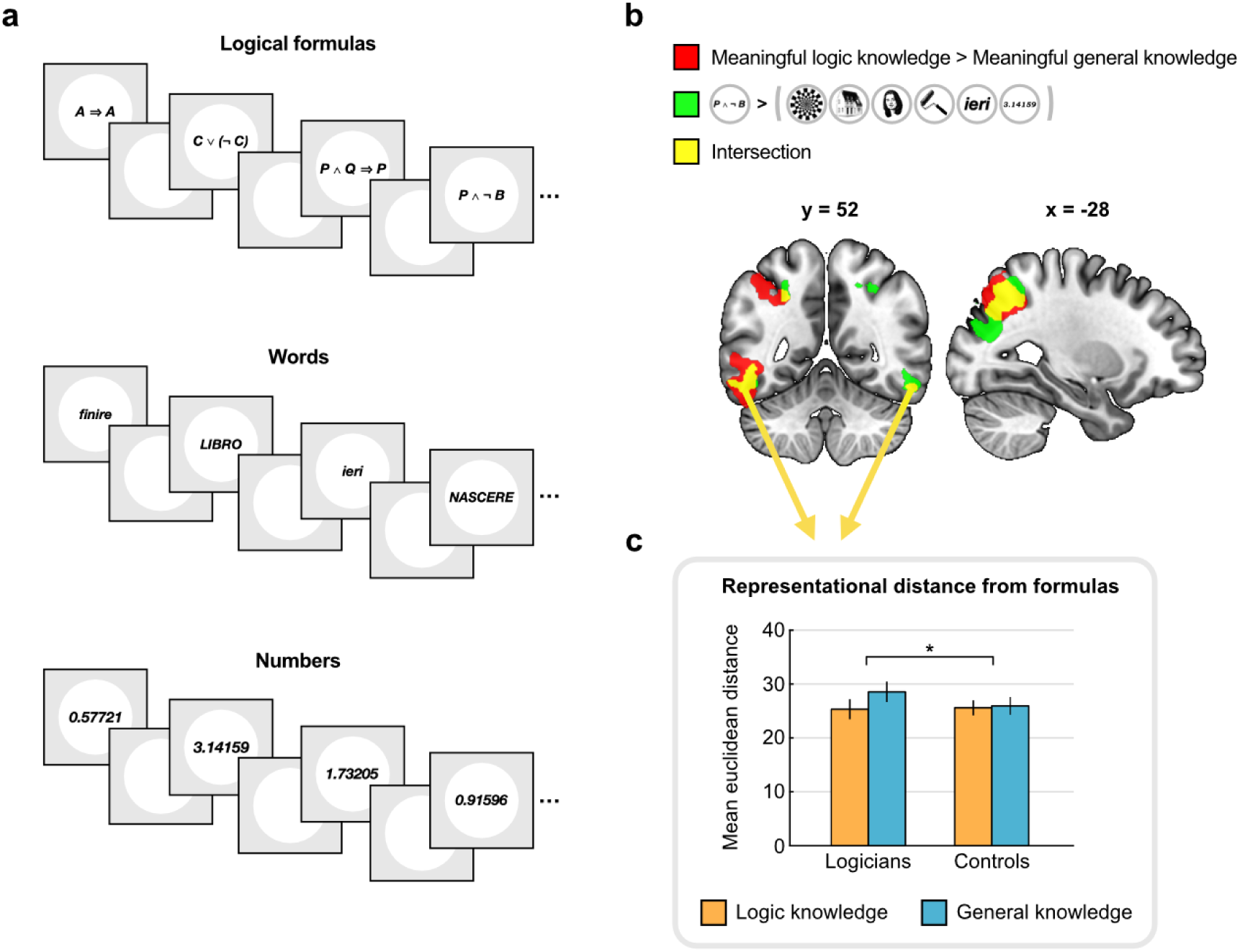
(a) Blocks of images of meaningful logic formulae, words, and decimal numbers, presented in an independent run, while participants performed a visual one-back task. (b) Whole-brain contrast maps of activation to meaningful logic statements vs. meaningful general knowledge (red), logic formulae vs. all other visual categories (green), and their intersection (yellow) in the group of logicians. (c) Euclidean distances between patterns of activation to formulae and those of logic statements (orange) vs. general knowledge statements (blue) in logicians and controls, computed within the ventral regions of intersection exhibited in (b).

In order to further characterize the response to logic formulae within the ventral visual cortex, first, we noted that in logicians only, the pITG response to written formulae overlapped with that evoked by logic spoken statements, bilaterally (Figure 5B). Quantification of this overlap using Dice coefficient confirmed substantial spatial correspondence between the logic written formulae and spoken statements (Dice = 0.10 +/− 0.037; 9 participants with p < 0.01 in hypergeometric tests; Z = 16.6; p < 0.001). We thus focused on these areas of overlap to characterize the patterns of response to formulae in relation to the other auditory and visual stimuli. Within these regions, at the individual level, we evaluated the neural representational similarity between the logic formulae and the other visual categories and between the logic formulae and the spoken statements. We first tested whether the representational distance between logic formulae and visual categories or spoken statements differed between groups. A mixed-model ANOVA with Group (logicians vs. controls) and Task (auditory statements vs. visual images) as factors, revealed for both groups, a higher proximity between logic formulae and other visual stimuli than between logic formulae and auditory statements (Main effect of Task: F(1,20) = 62.8, p < 0.001); no interaction with Group: F(1,20) = 0.73, p = 0.40). We then assessed whether formulae are representationally closer to logic-related statements than to general knowledge statements, and whether, as predicted, this effect was higher in logicians. A mixed-model ANOVA with Group (logicians vs. controls), Content (Formulae-Logic distance vs. Formulae-General Knowledge distance), and Hemisphere (left vs. right), showed main effects of Content (F(1,20) = 10.2, p = 0.005), and Hemisphere (F(1,20) = 196.7, p < 0.001), and a crucial interaction between Group and Content (F(1,20) = 6.65, p = 0.018). This indicates that the distributed patterns of activation to logic formulae are closer to logic statements in logicians than they are in controls (Figure 5C).

## Discussion

We investigated the neural representation of pure logic by examining how professional logicians evaluate statements concerning advanced logic concepts, compared to concepts of general knowledge. Four main findings emerged. First, evaluating statements about pure logic recruits a heavily left-lateralized brain network that includes the left intraparietal sulcus (IPS), left middle frontal gyrus (MFG), left dorsolateral prefrontal cortex, and the bilateral posterior inferior temporal cortex (pITG). Second, this network is fully segregated from the language network. Evidence for this dissociation comes from both the comparison between logic and general knowledge statements and the contrast between logic statements and an independent language localizer. Third, the logic network showed a large overlap with the left hemispheric component of the math-responsive network identified using an independent calculation localizer, but minimal with its right hemispheric counterparts. Fourth, written logic formulae activated ventral and parietal regions that partially overlapped with the logic-related network. Ventral activity in response to visual logic formulae slightly differed between logicians and controls in that its extent was larger and its representational pattern was more similar to that evoked by processing logic statements in logicians compared to controls, indicating an effect of expertise in the visual processing of written logic formula.

Historically, the conception of logic has shifted dramatically. Aristotle (384–322 BC) understood logic through the grammar of natural language, analyzing propositions as subject–predicate structures. Nineteenth-century developments in mathematics challenged this alignment: with the emergence of symbolic logic, most notably in Frege’s *Begriffsschrift* (1879), logical relations were detached from linguistic grammar and recast in symbolic notation. Logic then became a formal calculus designed to express laws of truth with the precision of mathematics^22^, and quickly shifted into the study of the structure and limits of formal symbolic systems that underlie our reasoning^1^. This historical trajectory illustrates the dual heritage of pure logic in both language and mathematics. The present findings suggest that, at the neural level, the conceptual representation of pure logic aligns more closely with its mathematical than to its linguistic foundations.

In fact, logic-related activations showed no spatial overlap with the classical language comprehension network identified by an independent localizer, and this despite the use of natural language as the common format for both logic and general knowledge statements, matched in word count, syntactic complexity, and delivery modality. Instead, language regions such as the temporal poles and anterior temporal cortex^23,24^ were primarily engaged when participants evaluated general knowledge statements. This dissociation indicates that the neural representation of logic concepts does not rely primarily on linguistic semantic systems, even when information is conveyed through spoken sentences. Our findings directly contradict theoretical positions that argue for the absence of any fundamental distinction between natural and formal languages^25^, as well as with the view that formal tools can be applied directly to the analysis of natural language^26^. However, our findings are consistent with previous work showing that mathematical knowledge can be accessed independently of language systems^7,8,27,28^ and extend this observation to the domain of pure logic. This dissociation has been further confirmed in all formal systems that have been studied such as arithmetic^29–31^, algebra^32,33^, or computer programming^2,3,34^. Taken together these findings raise the possibility that language processing alone—even at very high levels of sophistication—does not fully capture the neural and cognitive architecture underlying human logical reasoning (see also^35^). This may have important implications in the debate of whether artificial intelligence based on large language models can or cannot fully reproduce the abstract reasoning capacities characteristic of human cognition^36–38^.

Besides this striking dissociation with language, the network recruited during the comprehension of logic statements largely overlaps with activation elicited by mental arithmetic, and bears striking similarity with the math-responsive network (IPS, MFG, pITG) previously described in professional mathematicians^7,8^, and repeatedly implicated in the representation and manipulation of abstract symbolic structures in mathematics^30,32^. However, there was an intriguing exception: the overlap observed was strongly left-lateralized, while mathematical reasoning recruits bilateral fronto-parietal-temporal networks. Thus, the logic-responsive network seems to be mostly included in the math-responsive network. The absence of activity in the right parietal and frontal cortices during the evaluation of pure logic statements requires to be understood. One way in which pure logic differs from mathematics is that the former does not pertain to quantitative and/or spatial content, rather it focuses on the structures and relations, and refers to properties that are common across different symbolic systems used for reasoning. The reduced involvement of the right hemisphere observed in pure logic could therefore reflect the absence of quantity- or space-related computations. More generally, this observation raises the possibility that the math-responsive network contains partially distinct components: right-lateralized parietal circuits may support the representation of quantitative magnitudes and spatial relations^39–42^, whereas a more strongly left-lateralized subsystem may encode purely symbolic relational structures independently of magnitude or space. In this sense, these findings may resonate with longstanding debates in the philosophy of mathematics concerning the relation between mathematics and logic. Logicist programs such as those of Frege, Russell, and Whitehead attempted to reduce mathematics to logic, but ultimately encountered formal limitations internal to mathematical logic itself. Without bearing directly on those foundational questions, the present results suggest that mathematical cognition also relies on neurocognitive systems not fully shared with pure logical reasoning, particularly right-hemispheric parietal circuits implicated in quantitative and spatial representations. The partial dissociation between mathematical and logical processing may reflect not only a formal distinction between disciplines, but also differences in the neuro-cognitive architectures supporting them.

Our findings of a mostly left-lateralized fronto-parietal network for pure logic also converge with previous observations of left-dominant activity engaged during the execution of logical inferences such as Modus Ponens^12,13^ and during computer code comprehension tasks notably involving “if…then” functions^2^. This topographical alignment suggests that the brain represents concepts of pure logic like consistency or completeness by mobilizing the same neural machinery used to maintain and evaluate the internal coherence of a chain of thoughts within a formal system. Under this view, the left fronto-parietal network identified in our study could be seen as a general engine for formalization, which is subsequently applied to specific systems like arithmetic, propositional logic, or programming.

An additional insight into the nature of the representations engaged in this network comes from participants’ subjective reports. Logicians reported extremely low levels of mental imagery when evaluating logic statements. This contrasts with previous work in professional mathematicians, where participants reported relying on mental imagery when thinking about advanced math concepts. Furthermore, imagery ratings were correlated with activity in posterior inferior temporal cortex^7^, consistent with the idea that this region corresponds to “imagery node”^43^. Despite the absence of reported imagery in the present study, logic statements still recruited bilateral pITG. Since pure logic may represent an extreme form of abstraction, in which reasoning operates over the properties of formal symbolic systems themselves, rather than over quantities, spatial structures, or visualizable objects, this observation suggests that the involvement of the observed network is not determined by the use of visual mental imagery, but it instead reflects access to abstract symbolic knowledge. On the other hand, in light of the observation that people with aphantasia, who do not consciously report having mental imagery, consistently show activation in a closely related portion of the pITG^44^, it is also possible that, during logic processing, mental imagery is present but not available to conscious report.

Because logicians rated logic statements as more difficult and less well understood than general knowledge statements, and reported lower confidence when evaluating them, some readers could think that the observed activations reflect a high engagement of the so-called Multiple Demand Network, implicated in domain-general problem solving and cognitive control^45^. However, we can readily exclude such a difficulty account as logic-related activity persisted when difficulty was controlled for, both objectively (using subject’s accuracy) and subjectively (using their introspective difficulty ratings). These analyses demonstrate that the fronto-parietal-temporal network identified here cannot be explained by differences in task difficulty.

Building on previous findings that expertise can induce plastic changes in category-specific responses along the ventral visual pathway—for example in the form of increased activation for visual stimuli related to the domain of expertise (e.g., mathematical formulae in mathematicians, musical notation in musicians, or words in skilled readers)^7,20,21^—we investigated brain responses to a variety of visual stimuli, including logic formulae, in our two groups. We did not observe a major ventral stream reorganization in logicians compared to controls: the typical mosaic-like organization in response to the classical visual category^46^ did not differ across groups, and in both groups we identified a bilateral region whose activity was higher for written logic formula compared to all other visual categories. These results possibly indicate that the specific symbols composing the logical strings (letters, arrows, negation symbol, etc …) were equally visually familiar to both groups. Nonetheless, we did detect subtle group differences, indicating that the logicians’ ventral stream response to logic formulae differed from controls in two notable ways: first, in logicians, the spatial extent of the right hemisphere ventral cluster was larger; second its across-voxels pattern of activity was closer to that evoked during logic statement processing than during general knowledge statement processing. This suggests that formal training in logic subtly refines the visual ventral stream neural architecture towards mapping visual symbolic representations onto their underlying conceptual meaning.

To conclude, taken together the results from the current study indicate that pure logic, a discipline that straddles natural language and mathematics, relies on brain networks that are fully dissociated from the ones supporting natural language, and that are partially overlapping with mathematics. They point to an almost fully left lateralized cortical circuit that acts as a general engine for formalization, supporting purely abstract and symbolic thinking.

## Methods

### Participants

Twenty-four adult participants with normal or corrected-to-normal vision took part in the study. Twelve participants were professional logicians (ten men, two women; age: 42.6 ± 3.3 years; one left-handed), in that they were full-time researchers and/or university professors of logic and closely related fields (e.g., philosophy of logic and/or mathematical logic) operating within different departments: mathematics (n = 3), philosophy (n = 5) and computer science (n = 4). Twelve additional participants composed the control group (nine men, three women; age: 36.6 ± 2.1 years; one left-handed). They were also full-time researchers and/or university professors, but they had no knowledge of pure logic. Their disciplines were as follows: psychology (n = 6), ancient history (n = 1), language and translation (n = 2), chemistry (n = 1) and physics (n = 1). Age did not differ significantly between the two groups (t(22) = 1.53, p = 0.14), nor did education level, as all hold a PhD.

A sensitivity analysis (one-sample, one-tailed *t*-test, α = 0.001) using effect sizes reported in prior work investigating high-level math concepts in 15 professional mathematicians (Amalric & Dehaene, 2016) within the fronto-parietal-temporal math-responsive network (peak Cohen’s *d* ranging from 1.28 to 2.81), indicated high statistical power (> 0.90 for most regions) with samples of this size. Due to the rarity of professional logicians meeting our inclusion criteria, our final sample size equaled 12 participants, which provides adequate sensitivity to detect effects of comparable magnitude.

Prior to the experiment, participants gave written informed consent to participate. All procedures were approved by the Ethics Committee of the University of Trento.

### Protocol and Stimuli

Participants were scanned using fMRI while completing an auditory semantic evaluation task, a one-back visual category task, and a brief localizer of motor, visual, auditory, language, and mental calculation responses. After the scanning session, participants were presented with a questionnaire that probed, for each of the statements heard in the scanner, their degree of comprehension of the problem, the perceived difficulty, the certainty of their answer, the amount of mental imagery involved, and the type of reasoning involved (from memory retrieval to formal proof). While logicians evaluated all the statements they heard in the scanner, controls only evaluated the general knowledge ones.

#### Auditory semantic evaluation task

Participants performed an auditory semantic judgment task adapted from previous studies on mathematical knowledge^7,8,27,28^. Stimuli consisted of short spoken statements recorded in Italian, belonging to two main categories: pure logic and general knowledge. The pure logic statements were generated with the help of colleagues full professors in logic from various universities in Italy, and they referred to different formal systems (First-order logic, Peano Arithmetic, ZFC, Lambda calculus, Turing machines) and their properties (see the full list of sentences in the Supplementary Materials). The general knowledge statements pertained to different areas of knowledge (geography, history, biology, literature). Within each category, statements were further divided into three truth-value types: *true*, *false*, and *meaningless*. Meaningless statements were syntactically well-formed but semantically nonsensical combinations of words drawn from the meaningful ones. Half of the statements were declarative (e.g.*, “In ancient Greece, a citizen who was no longer able to pay their debts became a slave”; “The alphabet of a first-order language includes symbols for individual constants and for logical connectives”*) and the other half included informal inferential propositions (e.g., “*Since Italy lies on the Adriatic Sea, it has no coastline on the Pacific Ocean*”*; “If a first-order theory is categorical in the power of the continuum, then it is omega-stable”)*.

In total, the stimulus set comprised 96 statements (48 pure logic, 48 general knowledge controls), with 16 items per truth-value type. Statements were matched across categories for syntactic complexity and word count (logic: 16.7 ± 0.6 words; non-logic: 15.4 ± 0.4 words; t(94) = 1.87, p = 0.063). However, logic statements were slightly longer in duration than non-logical ones (logic: 5.4 ± 0.13 s; non-logic: 5.0 ± 0.13 s; t(94) = 2.28, p = 0.02). Each statement was presented once only.

The task was presented in a slow event-related fMRI design over 8 runs, each including 12 trials, such that each run included one exemplar of each subcategory [logic/general knowledge x propositions/inferences x true/false/meaningless]. Each trial followed the same structure (Figure 1A): An orange fixation cross first appeared on screen, and a 100-ms beep signaled the onset of the auditory statement, which was presented through MRI-compatible headphones. At the offset of the auditory statement, participants had 4 seconds to silently evaluate its truth value. Then, a second beep and a change of fixation cross to green signaled the response period. Participants were given 2 seconds to indicate whether the statement was true, false, or meaningless by pressing one of three buttons on an MRI-compatible response pad with their right hand. Each trial ended with a 7-second rest period during which the fixation cross returned to orange until the next trial. Participants completed three practice trials inside the MRI at the beginning of the exam to ensure task comprehension.

#### Visual run

To map brain activity along the ventral pathway, participants performed an additional visual run. It consisted of a one-back task including six different categories of stimuli: faces, houses, tools, words, numbers, and logic formulae, together with a control condition of circular checkerboards with greater retinotopic extent than the other stimuli. All images appeared as black on a white background (see Supplementary Materials, Figure S2). The task comprised 42 blocks (six per category), presented in randomized order. Within a block, 8 stimuli from the same category were shown for 300 ms each, separated by a 300 ms interstimulus interval. Participants were instructed to respond with their right index finger to consecutive stimulus repetitions, which occurred in 50% of the blocks.

#### Localizer Scan

A detailed description of the experimental protocol can be found in Pinel et al.’s original study^15^. In the present study, we adopted the same 5-minute fMRI localizer, with stimuli translated into Italian. In order to identify brain regions involved in language and mathematical processing, our analyses targeted two conditions: sentences (e.g., ‘In the city, taxis are easy to find’) and subtraction problems (e.g., ‘calculate thirteen minus seven’), presented both auditorily and visually, in different trials (Figure S3).

#### Post MRI questionnaire

After the fMRI session, participants completed a self-report questionnaire adapted from Amalric and Dehaene (2016). The questionnaire included all statements presented during the auditory semantic evaluation task, together with the response each participant had provided during scanning. Participants were instructed to base their ratings on their impressions during the scanning session rather than on any subsequent reflection.

For each statement, participants rated the following dimensions on a 7-point Likert scale: (1) comprehension (0 = no comprehension, 7 = complete comprehension), (2) certainty in their response (0 = random response, 7 = absolute certainty), (3) perceived difficulty (0 = very easy, 7 = very difficult), (4) type of reasoning involved (0 = no reasoning/factual knowledge, 7 = explicit reasoning), and (5) appeal to mental imagery (0 = none, 7 = very vivid mental images).

As in the main task, logicians evaluated both logic and general knowledge statements, whereas control participants evaluated only the general knowledge statements.

### Data collection and analysis

#### fMRI data acquisition and preprocessing

Magnetic resonance was performed on a 3-Tesla Siemens Prisma scanner using a 64-channel head/neck coil at the Center for Mind/Brain Sciences (CIMeC), University of Trento. Blood-oxygenation-level-dependent (BOLD) T2*-weighted functional images were acquired using a multiband echo-planar imaging (EPI) pulse sequence with the following parameters: Field of View (FoV) = 210mm; Voxel Size = 1.75 × 1.75 × 1.75mm; Number of slices: 75; Time Repetition (TR) = 2030 ms; Time Echo (TE) = 32 ms; Multiband acceleration (MB) factor = 3 and a flip angle of 65 °. In addition, a Multi-Echo MPRAGE GRAPPA sequence was used to acquire a T1-weighted structural image for each participant using the following parameters: parallel imaging acceleration factor (PI) = 2; FoV = 256mm; Voxel Size = 1 × 1 × 1mm; Matrix = 256 × 256 × 176, TR = 2530 ms, TEs = [1.69 ms, 3.55 ms, 5.41 ms, 7.27 ms] and a flip angle of 7 °.

The raw fMRI time series were preprocessed with a standard pipeline using SPM12, in Matlab R2019. Slice timing correction was based on the actual acquisition time of interleaved slices using the middle time point as reference. Functional images were then spatially realigned and motion corrected. Geometric distorsions were also corrected using the field maps. Structural images underwent segmentation into grey matter, white matter and cerebro-spinal fluid, and were coregistered to the mean functional image. Finally, both structural and functional images were normalized to the Montreal Neurological Institute (MNI) space and smoothed using a 4mm full-width-at-half-maximum (FWHM) isotropic Gaussian kernel. For both experimental tasks, preprocessed EPIs were high-pass filtered at 128 s and serial correlations were accounted for by using an autoregressive model AR(1).

#### Main fMRI Analysis

For the main experimental paradigm, as well as the localizer and the visual run, a general linear model (GLM) was computed for each participant and used to estimate subject-specific beta weights. Time series were modelled with the canonical SPM12 hemodynamic response function (HRF).

Within each of the 8 runs of the main experimental paradigm, a single boxcar regressor was created for each of the 12 presented statements. Each regressor modelled the activation associated with both the statement listening phase and the subsequent 4-s reflection period (pooled across). Regressors of non-interest included the six head motion parameters and two additional regressors modelling the activation related to the auditory alert signals and the button press responses. Subject-specific contrasts of interest were defined by comparing the activation evoked by each statement category relative to baseline. For the second level analysis, individual contrast images were smoothed with a Gaussian filter of 4mm FWHM, and entered into a second-level 12 × 2 whole-brain ANOVA with statement category as within-subject factor and group (logicians versus controls) as between-subject factor. To focus the analyses on positive activations, between-conditions contrasts (e.g., ‘meaningful logic > meaningful general knowledge’) were inclusively masked with one-sample t-tests against baseline level (e.g., ‘meaningful logic > baseline’) corrected at pvoxel < 0.05.

Time series of the localizer run were modelled by including 7 regressors, one for each experimental condition, as well as the six head motion parameters as regressors of noninterest. Button-press conditions were also modelled. For the present purposes, only two single-subject contrasts of interest were computed to isolate language processing ([sentence listening + sentence reading] > baseline) and mental calculation ([auditory mental subtraction + visual mental subtraction] > baseline).

For the visual run, time series were modelled by including one regressor for each of the 7 visual categories, as well as the six head motion parameters as regressors of noninterest. Button-press conditions were also modelled. For the second-level group analysis, individual contrast images for each of the experimental conditions relative to rest were smoothed with an isotropic Gaussian filter of 4 mm FWHM and entered into a second-level whole brain 6 x 2 ANOVA with visual category as within-subject factor and group as between-subject factor. Two subjects have been excluded from the analysis of the Visual Task for health-related reasons and malfunctions in the audiovisual setup.

All brain activations are reported using an uncorrected voxelwise threshold of p < 0.001 with a clusterwise false discovery rate (FDR) correction of pFDR < 0.05, accounting for multiple comparisons across the entire brain, and displayed on the MNI template.

#### Control for Difficulty Analyses

We conducted two kinds of control analyses: a group-level behavioral covariate analysis, and a within-participant analysis of subjective difficulty. The first control analysis aimed to regress out the potential effect of individual differences in task performance. To do so, the second-level model was re-computed with the inclusion of performance scores as a nuisance covariate. Specifically, each participant’s average accuracy was entered as a subject-level behavioral regressor, before evaluating the contrast of interest [meaningful logic > meaningful general knowledge].

The second control analysis, providing a more sensitive intra-individual control for difficulty, was performed based on participants’ subjective difficulty ratings in the post-MRI questionnaire. For each participant, meaningful statements were retrospectively sorted into two difficulty bins (easy and difficult) using a within-condition mean split: a statement was categorized as easy if its subjective difficulty rating was strictly below the subject’s own mean rating for that specific category and difficult if the rating was above the subject’s mean. A new first-level GLM was computed for each participant, with these sorted trials entered as separate regressors. The resulting subject-specific contrast images were then entered into a second-level whole-brain repeated-measures ANOVA with two within-subject factors: Condition (logic vs. general knowledge) and Difficulty (easy vs. difficult), before evaluating the contrast [easy logic > difficult non-logic].

#### Lateralization Index Analysis

In order to evaluate the amount of lateralization of the observed activation maps we used the Lateralization Index toolbox^47^. Regions of interest (ROIs) for both logic and language were constructed using the MarsBaR toolbox (https://marsbar-toolbox.github.io/) by creating 10mm-radius spheres centered on the peak coordinates of all significant clusters identified at group-level. To obtain a symmetric mask across hemispheres, ROIs were combined with their horizontally flipped counterparts, targeting the opposite hemisphere. These ROIs were then used as inclusive masks for single-subject unthresholded t-maps of interest for logic (meaningful logic > meaningful non-logic), language (sentence comprehension > calculation) and mathematical processing (calculation > sentence comprehension). Brain regions within 5mm off the hemispheric midline were used as an exclusive mask to remove possible flow artifacts.

For each participant, lateralization indices (LIs) were computed by means of a bootstrap algorithm^48^. This procedure implied the definition of 20 equally-sized t-values thresholds, ranging from 0 to the maximum t-value. At each threshold, 100 bootstrap samples were generated from the left and right ROIs, using a resampling ratio of k=0.25 (i.e., 25% of the data). This yielded a total of 10.000 lateralization indices, calculated using the standard formula [(Left-Right)/(Left + Right)] where Left corresponds to the number of suprathreshold voxels in the left hemisphere mask, and Right corresponds to the number of suprathreshold voxels in the right hemisphere mask. This procedure was repeated at each t-value threshold, for a final number of 200.000 LI combinations. To enhance reliability, we employed weighted estimates of LIs that assign greater importance to LI values obtained at higher t-value thresholds. The resulting LIs range from −1 to 1, where negative values indicate right-hemisphere dominance and positive values indicate left-hemisphere dominance. Absolute values reflect the strength of lateralization.

#### Spatial Colocalization Analysis

Spatial colocalizations between pairs of activation maps were quantified voxelwise using the Dice coefficient at the single-subject level: 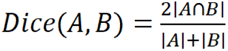, where 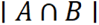 represents the number of voxels common to both maps, and |*A*| and |*B*| represent the number of suprathreshold voxels in each map. Dice values range from 0 (no overlap) to 1 (complete overlap). For each participant, unsmoothed single-subject contrast maps were thresholded at p<0.001 (uncorrected) and restricted to positively activated voxels within anatomical masks (a whole-brain gray-matter mask for the overlaps of logic or general knowledge statements with language processing or calculation, and a ventral mask anatomically defined as the bilateral inferior occipital, inferior temporal, and fusiform gyri, for the overlap between logic statements and logic formulae). To assess whether the observed spatial overlap for each participant was statistically significant, we modeled the voxel activation as a sampling process without replacement. For each participant, the exact probability of observing k or more overlapping voxels by chance was computed as P(X > k) from a cumulative hypergeometric distribution. This distribution was defined by the total number of voxels within the predefined mask (N), the number of active voxels in the first contrast (k), and the number of active voxels in the second contrast (n). Finally, to draw group-level inferences, individual p-values were aggregated using Stouffer’s Z-score method. Each subject-level p-value was transformed into a standard normal deviate (Z-score), and these were combined to compute a global Z-statistic. Logit-transformed dice coefficients were also entered into group-level analyses when appropriate.

#### Analysis of intensity and spatial extent of responses to formulae

To estimate the intensity spatial extent of formula-selective activation in the ventral visual stream, we followed the procedure adopted by Mongelli et al. in their study of the spatial extent of response to music scores^21^. We used the same anatomical mask of the left and right ventral cortex defined for the spatial colocalization analysis (inferior occipital gyrus, inferior temporal gyrus, and fusiform gyrus from AAL atlas). Within these masks, we identified the peak activation coordinates for the second-level contrast Symbols > Pictures pooling across both groups of logicians and controls (Left hemisphere: [−48, −56, −11], Right hemisphere: [55, −49, −14]). We then restricted each ventral mask around these peaks to a range of ±15 mm along the y-axis. For each participant, we identified the peak of the first-level contrast map of Formulae > Others, thresholded at p < .01 (uncorrected) and extracted the intensity of the activation as well as the extent of the cluster around that peak, within each restricted ventral mask (in number of voxels).

#### Representational Similarity Analysis

We started with the definition of functionally and anatomically constrained regions of interest: within the aforementioned anatomical masks of the left and right ventral cortex, we computed the intersection the binarized second-level logic activation map (Logic > General Knowledge, masked with Logic > Rest) and the binarized second-level formula activation map (Formulae > Others) in logicians. This procedure yielded two ROIs: left and right ventral intersection. For each participant (logicians and controls) and each ROI, we constructed a dataset concatenating two sets of beta estimates: beta estimates corresponding to meaningful logic and meaningful general knowledge statements from the main task, and beta estimates corresponding to the categories of houses, numbers, words, faces, tools, and formulae from the visual task. Beta patterns were first averaged across runs within each condition, then z-score normalized across voxels. Pairwise Euclidean distances between all condition pairs were computed using CoSMoMVPA. We ran two mixed-effect ANOVAs respectively assessing the effect of modality (auditory statements versus visual categories) and of statement semantic content (logic versus general knowledge) on composite values of distance to formulae, and its interaction with Group (logicians versus controls), adding participants as a random factor.

## Acknowledgements

This work was supported by a Marie Curie Global fellowship to M.A (Grant Agreement No. 839611 “NeuroMath: Acquisition of Mathematical Concepts in the Human Brain”), by the Italian Ministry of Education, University and Research under the PRIN 2022 programme to M.P. (Grant Agreement No. 2022EBC78W, “Sense of number vs. sense of quantity: modelling, neuroimaging, behaviour”), by the CARITRO foundation Post-Doc grant to P.A.M.M. This work was also supported by the European Research Council (ERC) under the European Union’s Horizon 2020 research and innovation programme (Grant Agreement No. 833504, SPANUMBRA, to G.V.). The authors wish to warmly thank professors Gabriele Lolli and professor Carlo Toffalori for their invaluable contribution in generating the statements of pure logic, and Professor Piergiorgio Odifreddi for discussions and help in recruiting logicians as well as all the colleagues who participated in the study as subjects.

## Supplementary Materials

### Supplementary figures

**Figure S1.**
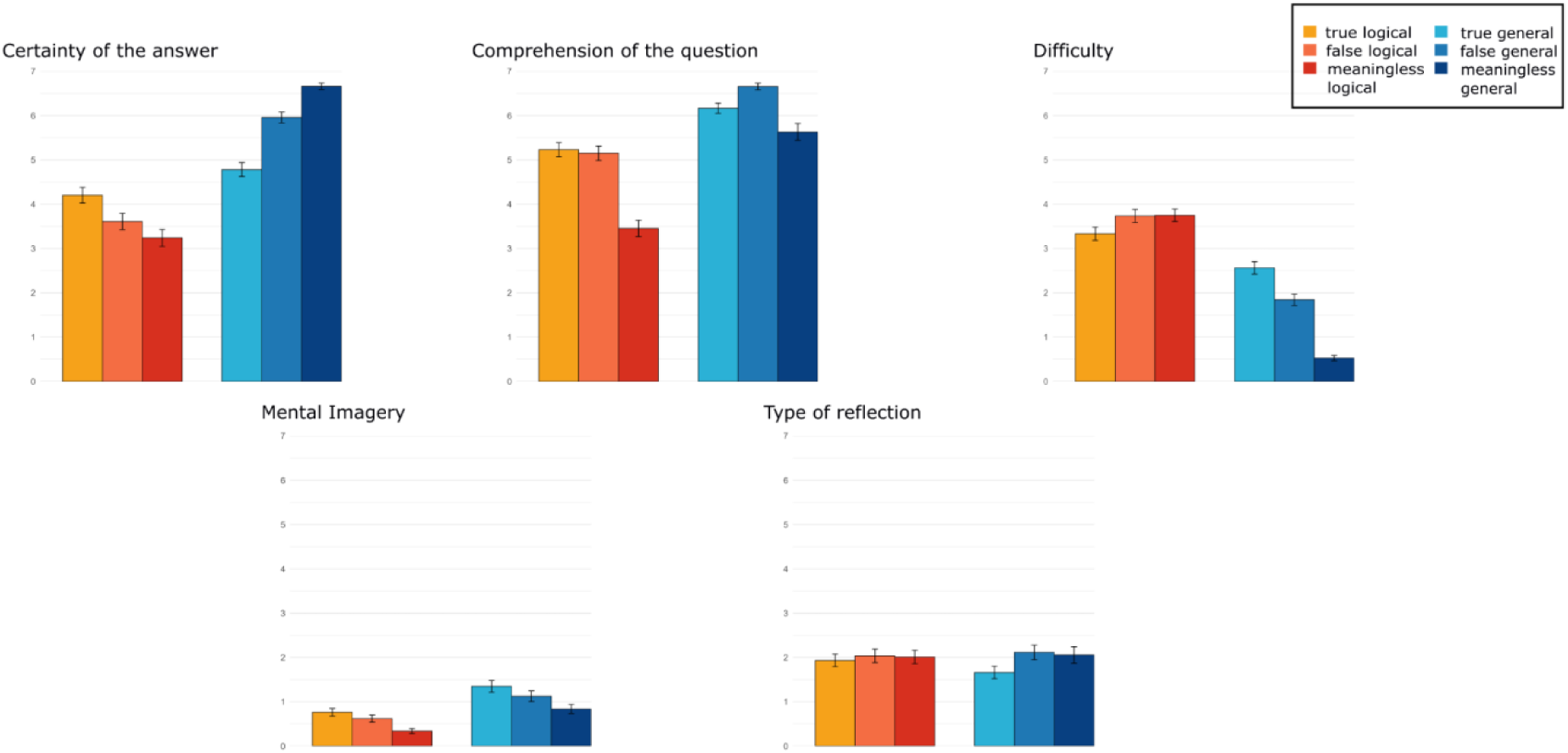
Logicians’ mean ratings in the post-MRI questionnaire

**Figure S2.**
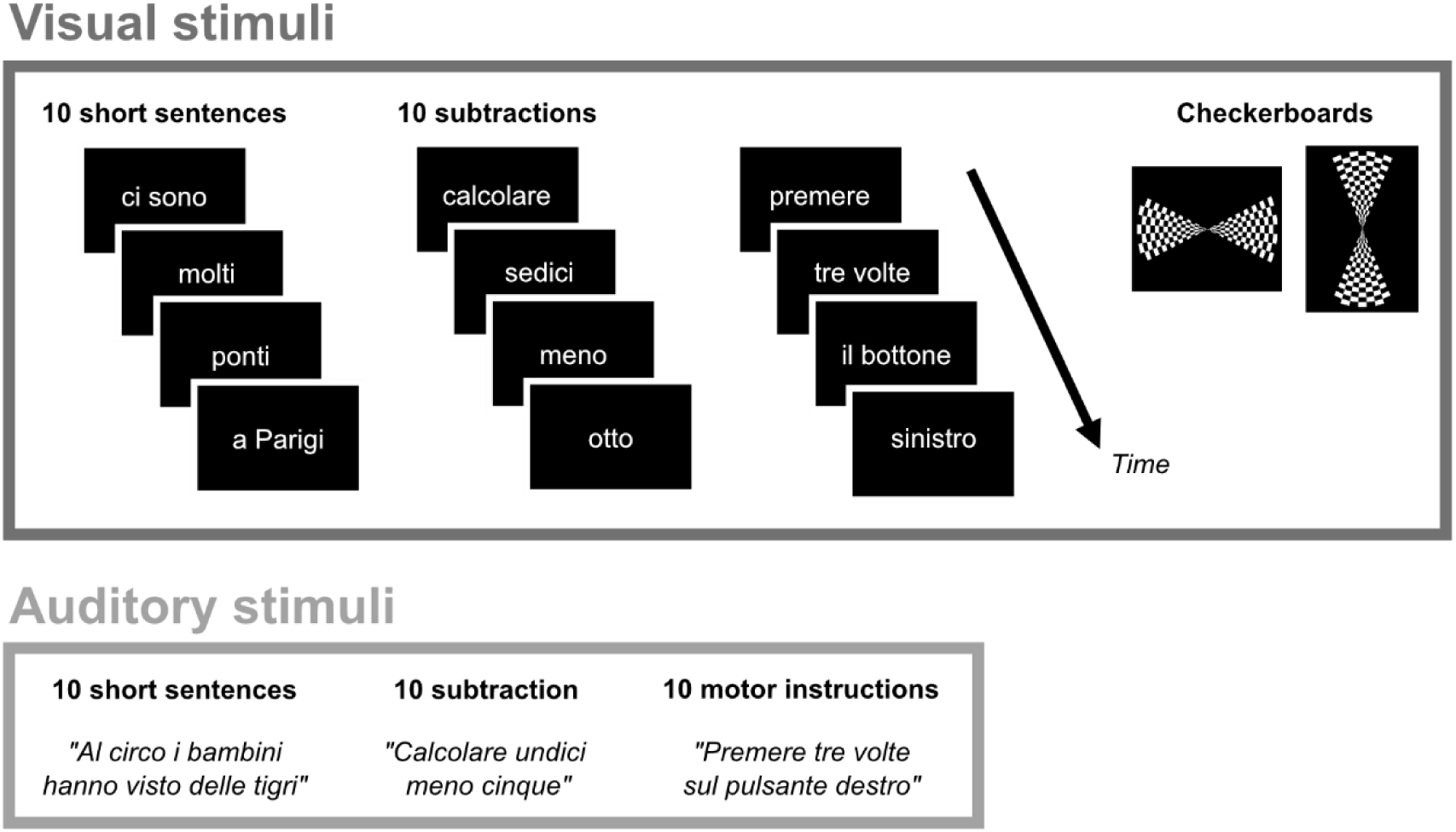
Examples of stimuli from the language and calculation localizer.

## References

1. The Development of Modern Logic. (Oxford University Press, 2009). doi:10.1093/acprof:oso/9780195137316.001.0001.

2. Liu, Y.-F., Kim, J., Wilson, C. & Bedny, M. Computer code comprehension shares neural resources with formal logical inference in the fronto-parietal network. eLife 9, e59340 (2020).

3. Liu (劉耘非), Y.-F. & Bedny, M. Learning to Program “Recycles” Preexisting Frontoparietal Population Codes of Logical Algorithms. J. Neurosci. 45, e0314252025 (2025).

4. Fedorenko, E., Behr, M. K. & Kanwisher, N. Functional specificity for high-level linguistic processing in the human brain. Proceedings of the National Academy of Sciences 108, 16428–16433 (2011).

5. Bedny, M., Pascual-Leone, A., Dodell-Feder, D., Fedorenko, E. & Saxe, R. Language processing in the occipital cortex of congenitally blind adults. PNAS 108, 4429–4434 (2011).

6. Dehaene, S., Piazza, M., Pinel, P. & Cohen, L. Three parietal circuits for number processing. Cognitive Neuropsychology 20, 487–506 (2003).

7. Amalric, M. & Dehaene, S. Origins of the brain networks for advanced mathematics in expert mathematicians. PNAS 201603205 (2016) doi:10.1073/pnas.1603205113.

8. Amalric, M. & Dehaene, S. A distinct cortical network for mathematical knowledge in the human brain. NeuroImage 189, 19–31 (2019).

9. Rodriguez-Moreno, D. & Hirsch, J. The dynamics of deductive reasoning: An fMRI investigation. Neuropsychologia 47, 949–961 (2009).

10. Monti, M. M., Parsons, L. M. & Osherson, D. N. The boundaries of language and thought in deductive inference. PNAS 106, 12554–12559 (2009).

11. Monti, M. M., Osherson, D. N., Martinez, M. J. & Parsons, L. M. Functional neuroanatomy of deductive inference: A language-independent distributed network. NeuroImage 37, 1005–1016 (2007).

12. Prado, J., Chadha, A. & Booth, J. R. The brain network for deductive reasoning: a quantitative meta-analysis of 28 neuroimaging studies. Journal of cognitive neuroscience 23, 3483–3497 (2011).

13. Goel, V. Anatomy of deductive reasoning. Trends in Cognitive Sciences 11, 435–441 (2007).

14. Wertheim, J. & Ragni, M. The Neurocognitive Correlates of Human Reasoning: A Meta-analysis of Conditional and Syllogistic Inferences. J Cogn Neurosci 32, 1061–1078 (2020).

15. Pinel, P. et al. Fast reproducible identification and large-scale databasing of individual functional cognitive networks. BMC Neuroscience 8, 91 (2007).

16. Cohen, L. & Dehaene, S. Specialization within the ventral stream: the case for the visual word form area. NeuroImage 22, 466–476 (2004).

17. Dehaene, S. & Cohen, L. The unique role of the visual word form area in reading. Trends in Cognitive Sciences 15, 254–262 (2011).

18. Yeo, D. J., Pollack, C., Merkley, R., Ansari, D. & Price, G. R. The “Inferior Temporal Numeral Area” distinguishes numerals from other character categories during passive viewing: A representational similarity analysis. NeuroImage 214, 116716 (2020).

19. Shum, J. et al. A Brain Area for Visual Numerals. Journal of Neuroscience 33, 6709–6715 (2013).

20. Dehaene, S. et al. How Learning to Read Changes the Cortical Networks for Vision and Language. Science 330, 1359–1364 (2010).

21. Mongelli, V. et al. Music and words in the visual cortex: The impact of musical expertise. Cortex 86, 260–274 (2017).

22. Pedriali, W. B. The Logical Significance of Assertion.

23. Fedorenko, E., Ivanova, A. A. & Regev, T. I. The language network as a natural kind within the broader landscape of the human brain. Nat. Rev. Neurosci. 25, 289–312 (2024).

24. Binder, J. R., Desai, R. H., Graves, W. W. & Conant, L. L. Where Is the Semantic System? A Critical Review and Meta-Analysis of 120 Functional Neuroimaging Studies. Cereb Cortex 19, 2767–2796 (2009).

25. Montague, R. Universal grammar. Theoria 36, 373–398 (1970).

26. Davidson, D. Inquiries into Truth and Interpretation: Philosophical Essays Volume 2. (Clarendon Press, 1984).

27. Moreno, A. et al. Languages of the brain: fMRI dissection of the amodal networks for language, mathematics, and social knowledge. 2025.12.19.695075 Preprint at 10.64898/2025.12.19.695075 (2025).

28. Amalric, M., Roveyaz, P. & Dehaene, S. Evaluating the impact of short educational videos on the cortical networks for mathematics. Proc. Natl. Acad. Sci. U.S.A. 120, e2213430120 (2023).

29. Nieder, A. & Dehaene, S. Representation of Number in the Brain. Annual Review of Neuroscience 32, 185–208 (2009).

30. Arsalidou, M. & Taylor, M. J. Is 2+2=4? Meta-analyses of brain areas needed for numbers and calculations. NeuroImage 54, 2382–2393 (2011).

31. Amalric, M. & Dehaene, S. Cortical circuits for mathematical knowledge: evidence for a major subdivision within the brain’s semantic networks. Phil. Trans. R. Soc. B 373, 20160515 (2018).

32. Maruyama, M., Pallier, C., Jobert, A., Sigman, M. & Dehaene, S. The cortical representation of simple mathematical expressions. NeuroImage 61, 1444–1460 (2012).

33. Monti, M. M., Parsons, L. M. & Osherson, D. N. Thought Beyond Language: Neural Dissociation of Algebra and Natural Language. Psychological Science 23, 914–922 (2012).

34. Ivanova, A. A., et al. Comprehension of Computer Code Relies Primarily on Domain-General Executive Brain Regions. http://biorxiv.org/lookup/doi/10.1101/2020.04.16.045732 (2020) doi:10.1101/2020.04.16.045732.

35. Kean, H. et al. Evidence from Formal Logical Reasoning Reveals that the Language of Thought is not Natural Language. 2025.07.26.666979 Preprint at 10.1101/2025.07.26.666979 (2026).

36. Mahowald, K. et al. Dissociating language and thought in large language models. Trends in Cognitive Sciences 28, 517–540 (2024).

37. Nezhurina, M., Cipolina-Kun, L., Cherti, M. & Jitsev, J. Alice in Wonderland: Simple Tasks Showing Complete Reasoning Breakdown in State-Of-the-Art Large Language Models. Preprint at 10.48550/arXiv.2406.02061 (2025).

38. Shojaee, P. et al. The Illusion of Thinking: Understanding the Strengths and Limitations of Reasoning Models via the Lens of Problem Complexity. Advances in Neural Information Processing Systems 38, 108018–108059 (2026).

39. Warrington, E. K. & James, M. Tachistoscopic number estimation in patients with unilateral cerebral lesions. J Neurol Neurosurg Psychiatry 30, 468–474 (1967).

40. Kinsbourne, M. Hemi-neglect and hemisphere rivalry. Adv Neurol 18, 41–49 (1977).

41. Kimura, D. Dual functional asymmetry of the brain in visual perception. Neuropsychologia 4, 275–285 (1966).

42. Mesulam, M.-M. A cortical network for directed attention and unilateral neglect. Annals of Neurology 10, 309–325 (1981).

43. Bartolomeo, P., Liu, J. & Spagna, A. The Fusiform Imagery Node: Where vision meets concepts in the left temporal lobe. Neuropsychologia 224, 109398 (2026).

44. Liu, J. et al. Visual mental imagery in typical imagers and in aphantasia: A millimeter-scale 7-T fMRI study. Cortex 185, 113–132 (2025).

45. Duncan, J. The multiple-demand (MD) system of the primate brain: mental programs for intelligent behaviour. Trends in Cognitive Sciences 14, 172–179 (2010).

46. Kanwisher, N. & Dilks, D. The Functional organization of the ventral visual pathway in humans. in The New Visual Neuroscience (2013).

47. Wilke, M. & Lidzba, K. LI-tool: A new toolbox to assess lateralization in functional MR-data. Journal of Neuroscience Methods 163, 128–136 (2007).

48. Wilke, M. & Schmithorst, V. J. A combined bootstrap/histogram analysis approach for computing a lateralization index from neuroimaging data. NeuroImage 33, 522–530 (2006).

